# drMD: Molecular Dynamics for Experimentalists

**DOI:** 10.1101/2024.10.29.620839

**Authors:** Eugene Shrimpton-Phoenix, Evangelia Notari, Christopher W. Wood

**Affiliations:** School of Biological Sciences, University of Edinburgh, Roger Land Building, Edinburgh EH9 3FF; School of Chemistry, University of Edinburgh, Joseph Black Building, Edinburgh, EH9 3FJ

## Abstract

Molecular dynamics (MD) simulations can be used by protein scientists to investigate a wide array of biologically relevant properties such as the effects of mutations on a protein’s structure and activity, or probing intermolecular interactions with small molecule substrates or other macromolecules. Within the world of computational structural biology, several programs have become popular for running these simulations, but each of these programs requires a significant time investment from the researcher to run even simple simulations. Even after learning how to run and analyse simulations, many elements remain a “black box.” This greatly limits the accessibility of molecular dynamics simulations for non-experts.

Here we present drMD, an automated pipeline for running molecular dynamics simulations using the OpenMM molecular mechanics toolkit. We have created drMD with non-experts in computational biology in mind. The drMD codebase has several functions that automatically handle routine procedures associated with running molecular dynamics simulations. This greatly reduces the expertise required to run MD simulations. We have also introduced a series of quality-of-life features to make the process of running MD simulations both easier and more pleasant. Finally, drMD explains the steps it is taking interactively and, where useful, provides relevant references so the user can learn more. All these features make drMD an effective tool for learning molecular dynamics while running publication-quality simulations.

drMD is open source and can be found on GitHub: https://github.com/wells-wood-research/drMD.

## Introduction

Protein scientists are confronted with a pair of problems: proteins are far too small to see and constantly moving. This makes the process of understanding protein structures at the atomic level challenging. Experimental techniques can be used to probe protein structure and dynamics, but these often paint an incomplete picture. For example, X-ray crystallography can be used to investigate protein structures at the atomic scale, but provides very little information on protein dynamics [1]. Conversely, nuclear magnetic resonance (NMR) can be used to study protein dynamics. However the interpretation of the raw data from NMR experiments is far from reliable, which prevents the technique from becoming standard practice for protein scientists [2]. Molecular dynamics (MD) uses physics-based methods to simulate the real-time motions of atoms in a system at the femtosecond timescale. All-atom simulations are a complementary tool to these experimental methods and can provide more information on protein dynamics.

MD has been used to investigate a wide range of biologically important properties, such as the role of single-point mutations on enzyme structure [3,4] or activity [5,6] and the quantitative determination of binding energies for both protein-ligand [7,8] and protein - protein complexes [9]. Hybrid quantum mechanical / molecular mechanical (QM/MM) methods commonly use MD simulations as sampling methods [10], which allows for several different starting geometries to be used when calculating potential energy surfaces of reaction pathways. With the recent advent of deep learning-based structure prediction methods [11–13], MD methods have found a new use; energy minimisation steps using MD methods are commonly applied to predicted structures to remove unphysical steric clashes [14].

There are currently many software packages available to run MD simulations. These include OpenMM [15], GROMACS [16], NAMD [17], AMBER [18] and, CHARMM [19]. Broadly, each of these software packages gives researchers access to the same kinds of simulations, with comparable levels of accuracy and speed [20]. Each package provides slightly different collections of specialised features such as enhanced sampling or coarse-grained simulations, although this is rarely relevant to a newcomer to the field. The greatest difference between these packages is their ease-of-use. Several attempts have been made to make running MD simulations more user-friendly. CHARMM-GUI [21] has a web-based graphical user interface that makes much of the user input process more intuitive for those with little coding experience. Alternatively, OpenMM [15] has been created to be fully Python compatible and can be run from within custom Python scripts. Neither of these approaches however serve to simplify the actual decision-making process that goes into the set-up of MD simulations.

Here we present drMD (available at https://github.com/wells-wood-research/drMD,), an automated pipeline (see Figure 1) for running molecular dynamics simulations using the OpenMM MD toolkit [15] with the AMBER forcefield [18]. The purpose of drMD is to greatly reduce the complexity of setting up MD simulations, thus reducing the expertise required to run them. It is our hope that this will encourage non-computational protein researchers to use MD simulations to further their research goals. To this end, we have developed several quality-of-life features absent from other molecular dynamics packages (for detailed list of these features, consult our supplementary information). Unlike other MD packages, drMD accepts a single configuration file as its argument. This allows for all the important decisions to be made at once and to be stored in the same file. We believe that this provides clarity to the process of setting up simulations and is vital for reproducibility.

**Figure 1:**
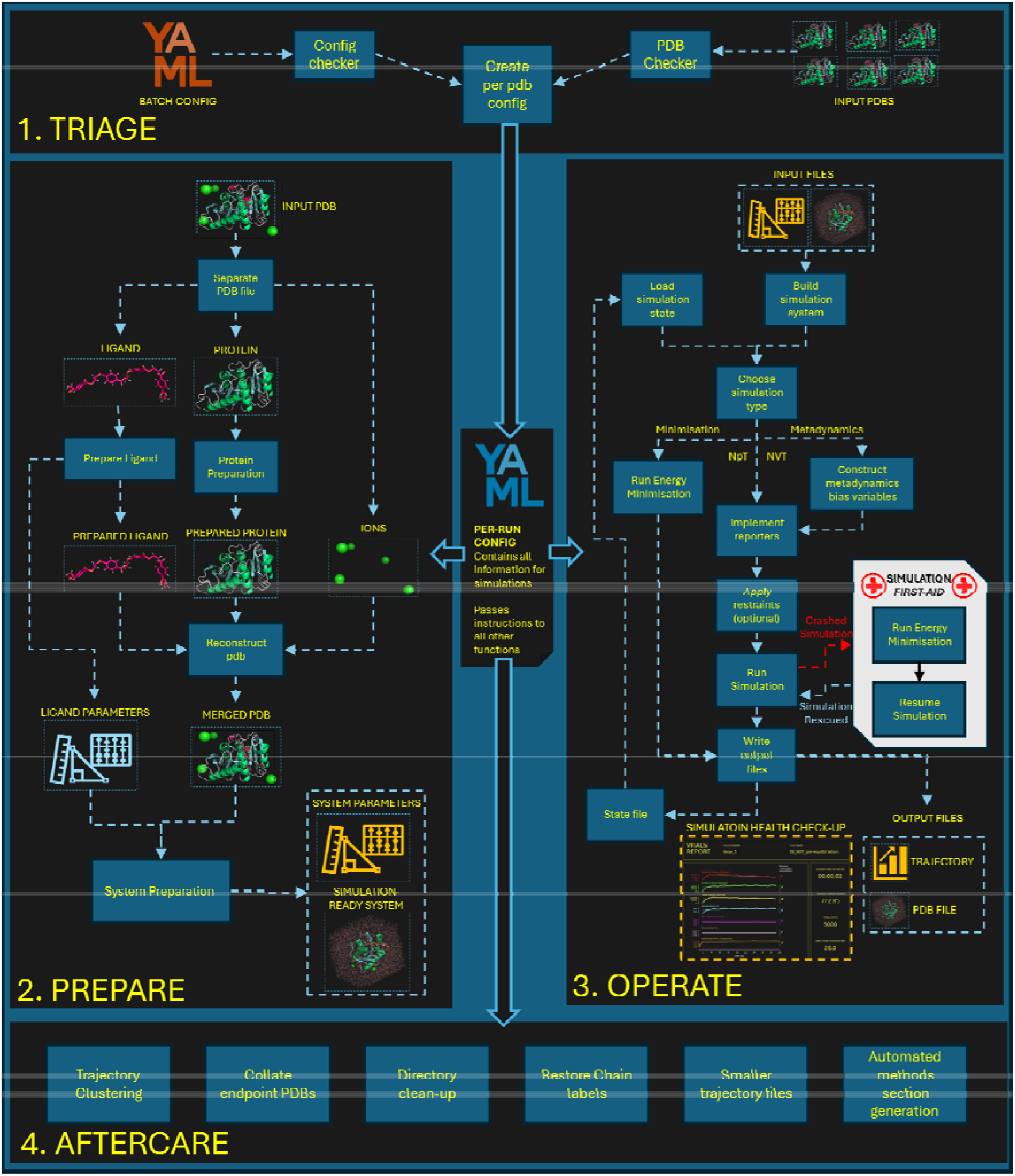
A visual representation of the drMD workflow. drMD first performs “triage” steps where input files are validated. Next, preparation steps are performed to generate input files for simulations, which are then performed using OpenMM. Once all simulations have been performed, some optional “aftercare” steps can be performed.

drMD will produce a “vitals report” after running each simulation (see figure SI II), producing a PDF that can be used to assess whether a simulation has converged with a glance. drMD also has an optional feature that can be used to “rescue” simulations that have failed due to some numerical error. Combined with many other quality-of-life features, drMD is a powerful and user-friendly way for newcomers to MD to run simulations.

## Results and Discussion

### The drMD configuration file

When drMD is run from the command line, it takes a single YAML file as an input argument. Within this configuration file, the user can define all the details required to run their simulations of interest. This information includes paths to input and output files, as well as details on their system’s hardware. This configuration file can be used to run multiple sequential MD simulations with each one using the previous simulation’s final geometry as its input. drMD can also apply a protocol described in a configuration file to several input structures. These features greatly streamline the process of running MD simulations.

The drMD codebase allows for all of the most commonly used MD simulation types to be performed. Simulations under both the *canonical* (a.k.a. NVT) and the isothermal*-isobaric* (a.k.a. NpT) ensembles may be performed with drMD, as well as simple energy minimisation optimisations. We have also implemented metadynamics simulations into the drMD codebase. This will allow users to perform enhanced sampling, which can be invaluable in exploring complex energy landscapes.

We have implemented distance, angle, torsion and position restraints into drMD, which allows the user to apply additional forces to atoms in the simulation, and is often central to equilibration procedures [22]. We have implemented a robust and flexible atom selection method in our configuration files. This method is reused throughout the configuration file to specify restraints, create bias variables for metadynamics, and to specify atoms to be included in the output files.

Having all the input information in a single file contributes to the reproducibility of the simulations run using drMD. A configuration file can be provided in the supplemental information section of a paper, which can be then used by anyone to reproduce the original simulation. We provide detailed instructions on how to format this configuration file in our GitHub readme file. We have also provided some supplemental instructions that can guide a researcher new to the field in what sorts of simulations they want to run. Should an incorrectly formatted configuration file be submitted to drMD, an exhaustive configuration file checker algorithm will provide the user with guidance on how to correct any issues with the file.

### Automation of preparation and simulation steps in drMD

A major challenge in the field of MD is the parametrisation of small, non-protein molecules. The presence of an unparameterised molecule will cause a simulation to fail. drMD automates this process using parmchk from AMBER [18] and OpenBabel [23].

For simulations to have biologically meaningful results, all molecules present must be in their correct protonation state. drMD automates protein protonation using pdb2pqr [24,25] and ProPKA [26] and the protonation of small molecules with OpenBabel [23].

drMD also automates the serialisation of simulation steps. This allows for a long series of equilibration steps to be performed prior to the production simulation, without constant intervention on the part of the user.

### Quality-of-life features in drMD

Many quality-of-life features have been incorporated into the codebase of drMD, each with the aim of making the process of running MD simulations as frictionless as possible. The following features have been implemented:

- Pre-built configuration files
- Extensive configuration file validation
- Simple input PDB checker
- Parallelisation of simulations
- Real-time updates of simulation progress
- Retention of chain and residue information
- Simulation “vitals reports”
- Automated methods-section generation
- Trajectory clustering
- Collation of simulation end-points
- Automated directory clean-up
- Implementation of logging
- “First-Aid” protocol for “rescuing” crashed simulations

For a more detailed explanation of each of these quality-of-life features, see our supplementary information.

We note that drMD will not be an appropriate tool for users who wish to run certain advanced simulation types such as funnel metadynamics to assess ligand binding energies [7]. drMD does not currently support the ability to run simulations on more esoteric structures such as proteins with non-natural amino acids, post-translational modifications, or organometallic ligands. For those wishing to perform such simulations, rather than starting from scratch, the open-source licence of drMD makes it ideal for the user to modify it to add any required functionalities.

### Example simulations with drMD

To demonstrate the utility of drMD, we chose to replicate a series of MD simulations performed by Lemeira *et al*. on the IsPETase enzyme [27].

Using the docking software GNINA [28], we were able to qualitatively reproduce the binding pose of the PET tetramer reported in the work of Lemeira *et al*. [27]. Using our drMD pipeline, we have reproduced the simulations performed by Lemeira *et al*. [27]. Simulations were performed upon IsPETase in its apo form (system I) and its holo form, bound to a PET tetramer (system II). We recreated the equilibration and simulation protocol from the original paper with some difficulty. This was not due to any technical issues, but rather due to ambiguity in how the protocol was presented in the original paper. With drMD, the entire protocol is contained in a single YAML configuration file. If one wished to reproduce this work again, or apply the simulation protocol to a different system, this configuration file could be re-used, making the process trivial.

Once the solvation and equilibration protocol (detailed in the Supplementary Information) had been performed, a quick visual inspection drMD’s simulation vitals report revealed that the system had reached convergence by the end of the equilibration simulation step (see figures SI III and SI IV). A similar check of the vitals report corresponding to the production MD step showed that these simulations were performed under equilibrium conditions (see figures SI III and SI IV).

Analysis of the trajectory of systems I and II showed that our methods produced qualitatively similar MD simulations to those of Lemeira *et al*. [27]. In general, no large-scale motions were observed in either system. We performed a comparison of root-mean-squared fluctuations (RMSF) of alpha-carbons in systems I and II (see Figure 3). The majority of residues in system II have lower RMSF values (i.e., they are moving less during the simulation) than their corresponding residues in system I. This suggests that the presence of the PET-tetramer has a stabilising effect on IsPETase. This effect is more pronounced within the active site. Residues that are most stabilised by the presence of the PET-tetramer reside on loops 4, 5, 7, 9, 10 and 13 (see Figure 2). Each of these loops are located at the active site of the enzyme. An increase in RMSF was observed in system II on loop 3 as well as the N-terminal and C-terminal loops 1 and 16 respectively (see Figure 2). These results are in broad agreement with those of Lameria *et al*.

**Figure 2:**
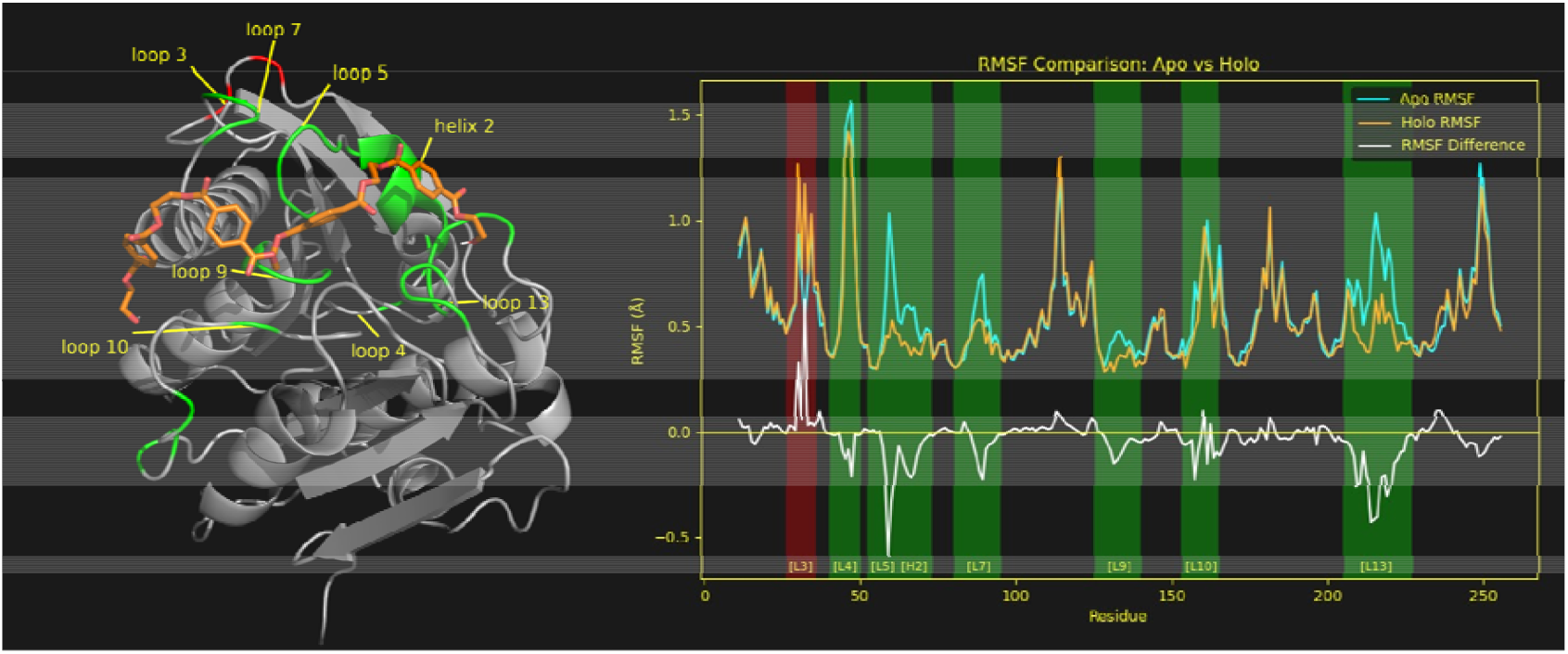
**Left** shows the structure of IsPETase with a PET-tetramer bound to its active site (shown orange). The cartoon representation is coloured by the difference of the RMSF values between systems I and II, with green representing a significant reduction in RMSF upon PET-tetramer binding and red representing a significant increase in RMSF. **Right** shows traces for RMSF per residue for System I (cyan) and II (orange), and the difference in RMSF between the two systems (white). Regions that have been stabilised and destabilised upon PET binding are labelled in green and red respectively. Note that, for clarity, the values for the C and N termini have been omitted from the plot.

## Conclusions

In this article, we have presented drMD, a python-based software that can be used to run MD simulations on proteins and protein-ligand complexes. drMD is a fully automated pipeline that handles system preparation, performs sequential simulation steps, and provides postprocessing for simulation output files. This includes the handling of difficult steps such as ligand parameterisation, restraints and enhanced sampling.

Much of the design behind drMD has been tailored towards protein scientists without much computational expertise. Biomolecular simulations have the potential to be an invaluable tool for experimentalists as they can be used to rationalise wet-lab results and to guide future research. Through drMD’s many quality-of-life features, as well as our emphasis on reproducibility, we aim to further the field of protein science by making high-quality MD simulations more accessible.

## Acknowledgements

We thank members of the Wells Wood Research Group for testing and feedback on the drMD. Eugene Shrimpton-Phoenix and Christopher W. Wood are supported by a BBSRC sLOLA award (BB/X003027/1). Evangelia Notari is supported by a PhD studentship from the UKRI funded EastBio Doctoral Training Partnership programme.

