## Supplementary Information for "drMD: Molecular Dynamics for Experimentalists"

### drMD’s quality-of-life features

drMD contains a large amount of features that aim to make the process of running biomolecular simulations smoother and more pleasant. This section contains more exhaustive list of these features than could be reasonably included in the main text of the paper:

#### Extensive config checker

drMD takes a single configuration file as its input. We have made great effort to make the formatting of this configuration file intuitive and have provided extensive instructions in our GitHub repository (https://github.com/wells-wood-research/drMD). It is however likely that users will occasionally submit configuration files that are missing key information or are formatted improperly. Instead of throwing a generic error message and forcing the user to work out what is wrong on their own, drMD will inform the user the exact section of the configuration file that has caused the problem and provide helpful tips on how to fix it. We hope that this feature will make drMD far more user-friendly than a lot of other command-line applications in the field.

#### Simple PDB checker

There are myriad ways that an input PDB file can cause the various pieces of software that drMD uses to throw errors. Before any simulations are run, drMD will look through all the input PDB files and test them for the following common problems:

- Discontinuous chains
- Residues with multiple conformers
- Residues with missing atoms
- Non-natural amino acids
- Organometallic ligands

Many of these problems can be fixed using the OpenMM’s PDBFixer [1]. This feature is optional and can be disabled by setting the **skipPdbTriage** entry in the config file to False.

#### Parallelisation of simulations

If the user provides an input of more than one in the **paralellCpus** field of the config file, drMD will run their simulations in parallel. This is presented with an attractive series of loading bars in order to let the user know how long the simulations will take.

#### Real-time updates of simulation progress

While simulations are running, drMD provides the user with information on how long the simulations will take, as well as the average speed of the simulation. This information is displayed on the command line and is updated every few seconds. This is useful as it allows the user to better plan their time.

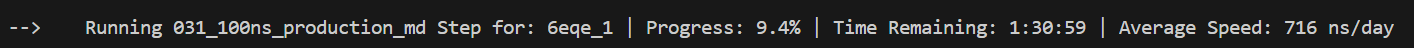

Figure SI I: An example of the real-time time updates that drMD provides the user as their simulations are running.

#### Retention of chain and residue information

By default, OpenMM discards chain information and sets all chains to start with residue one. This can be extremely frustrating as one often finds that residues in a crystal structure no longer correspond with those in MD output files. drMD implements a simple function at the end of each simulation step that reinstates chain information and resets residue numbers to their original values.

#### Simulation vitals reports

drMD gathers information on simulations as they are running using a combination of OpenMM’s built-in reporters [1] and reporters from the MDAnalysis package [2,3]. On their own, these reporters simply create CSV files. drMD automatically plots the contents of these reporters and collates them into a “vitals report” PDF file (see figure SI II below). A simple gradient-based algorithm is then performed to assess whether the following key simulation properties are at equilibrium during the simulation:

- Backbone RMSD
- Total Energy
- Kinetic Energy
- Potential Energy
- Simulation Box Volume
- Simulation Box Density
- Temperature

If any of these properties are not at equilibrium, we advise against performing further analysis. Instead, further equilibration simulations are most likely required.

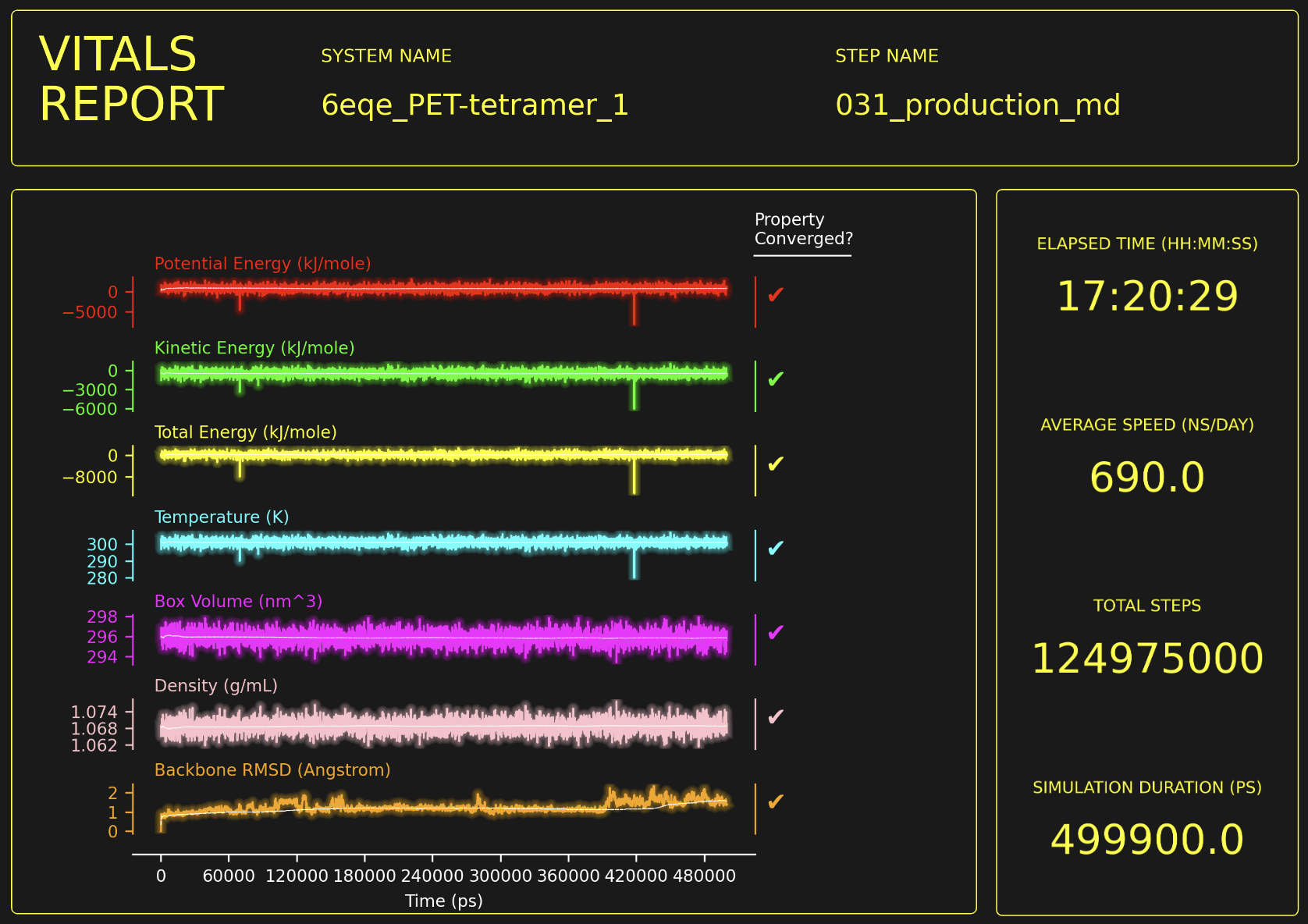

Figure SI II: An example of a simulation vitals report. For a more detailed explanation of how to interpret these reports, see the **Simulation Health Checks** section later in this document.

#### Automated methods-section generation

For non-experts, writing a methods section for MD simulations can be rather daunting. Out of the several steps that are required to set up and run a simulation, it is often difficult to tell which steps need to be included in the methods section of a paper or a thesis. It is also laborious to collect all of the relevant citations for this section. After drMD has finished running simulations, it will gather the relevant data by looking through all the per-run configuration files as well as all of the preparation steps. It will then programmatically generate a methods section file (located in the 00_AutoMethods directory in the user’s specified **outputDir**). This methods section contains every detail that needs to be included, with accompanying citation DOI links. An example of one of these automated methods sections is provided later in the section **Automatically generated methods section for molecular dynamics protocol**.

As this methods section is generated by a hard-coded algorithm, it may read as robotic and overly verbose. We recommend that the user manually reformats the methods section for publication. We also recommend that the automatically generated methods section is supplied as-is in a publication’s supplementary information, as this ensures that the protocol can be replicated exactly.

#### Trajectory clustering

Long MD simulations create a lot of data and trajectory files can be very large. Often, a user will simply want to see the most interesting snapshots of a simulation, rather than having to manually look through each frame of the simulation. To aid this, drMD implements an optional trajectory clustering step. This step performs k-means clustering upon the Cartesian coordinates of a simulation trajectory using a combination of MDAnalysis [2,3] and scikit-learn [4]. This generates a user-defined amount of PDB files, each representing a cluster of structures. By using this feature of drMD, the user can get a quick idea of the range of poses that their system adopts throughout their simulation.

#### Collation of simulation end-points

Once all of the simulations have completed, drMD can optionally collect all of the resulting end-points of simulations into appropriate directories, with appropriate file names. This is especially useful if it is used to perform many energy minimisation calculations. In this case, the final geometry of each system is the only thing the user is interested in. As such, collating these geometries into one directory is very useful.

#### Automated directory clean-up

Running a series of MD simulations can create a lot of files, some of which can be rather large. drMD provides the option to delete all simulation directories, which is useful in combination with trajectory clustering or end-point collation as described above. Alternatively, drMD provides the option to delete directories associated with a particular simulation step. Whilst this feature can save the user a lot of storage, we recommend caution when using it.

#### Implementation of logging

At each step throughout drMD’s preparation and simulation processes, a log of what is happening is written to file. These logs can be useful in identifying root causes for errors. Log files are written to the 00_drMD_logs directory within the user specified **outputDir** directory.

#### Rescuing crashed simulations with “First Aid”

Sometimes MD simulations “explode” due to numerical errors. This most commonly occurs when an energy of an interaction is evaluated as infinite, for example when a pair of atoms overlap. We have implemented a “First Aid” protocol into drMD: If a simulation fails due to a numerical error, drMD will perform an energy minimisation starting from the last useable geometry from the exploded simulation. Once a minimised structure has been obtained, drMD will attempt to run the original simulation from the timestep where the explosion took place. This process can be performed multiple times (controlled by the **firstAidMaxRetries** parameter in the config file) and can be used to perform simulations that would usually be impossible.

We caution against relying on this feature too much. If you simulation keeps exploding, consider reducing the **timestep** or **temperature** parameters of the simulation.

### Methods section for demonstration simulations

#### Molecular docking protocol

To generate the binding pose of the PET-tetramer (HE-(MHET)_4_), we performed molecular docking using the program GNINA [5] (a GPU accelerated version of vina [6] that uses convolutional neural networks). We used a docking box with dimensions 40 x 40 x 15 Å with its central coordinate defined by the coordinate of the catalytic cysteine’s gamma oxygen atom (Ser160 - OG). We used an exhaustiveness parameter of 16 and used the seed 42. In contrast to the methods we wished to replicate, we performed this docking simulation without any flexible residues in accordance to the recommendations on the GNINA GitHub. We proceeded with the binding pose with the greatest GNINA CNN score.

#### Automatically generated methods section for molecular dynamics protocol

**Molecular Dynamics Protocol**

**This methods section was automatically generated by drMD [Ref.** [**drMD**](https://doi.org/PLACEHOLDER)**].**

This document contains all the information needed to recreate your simulations.

Feel free to use this as the basis for your methods section in your papers, thesis, etc.

**Ligand Preparation and Parameterisation**

Ligand parameter generation was performed using the following procedure: All ligands were protonated using OpenBabel [Ref. [obabel](https://doi.org/10.1186/1758-2946-3-33" \o "https://doi.org/10.1186/1758-2946-3-33)], newly added hydrogen atoms were then renamed to ensure compatibility with the AMBER forcefield. Partial charges of all ligands were calculated, and atom types were assigned using antechamber [Ref. [antechamber(1)](https://doi.org/10.1002/jcc.20035), and [antechamber(2)](https://doi.org/10.1016/j.jmgm.2005.12.005)], and the parameters for the ligands were generated using parmchk [Ref. [parmchk(1)](https://doi.org/10.1002/jcc.20035" \o "https://doi.org/10.1002/jcc.20035), and [parmchk(2)](https://doi.org/10.1016/j.jmgm.2005.12.005" \o "https://doi.org/10.1016/j.jmgm.2005.12.005)].

**Protein Preparation**

All proteins were protonated using software pdb2pqr [Ref. [pdb2pqr(1)](https://doi.org/10.1093/nar/gkm276), and [pdb2pqr(2)](https://doi.org/10.1093/nar/gkh381)] which uses ProPKA to calculate per-residue proton affinities [Ref. [propka(1)](https://doi.org/10.1021/ct200133y" \o "https://doi.org/10.1021/ct200133y), and [propka(2)](https://doi.org/10.1021/ct100578z" \o "https://doi.org/10.1021/ct100578z)]. Proteins were protonated using the following pH: 7.4. This process also automatically creates disulfide bonds as appropriate.

**Solvation and Charge Balancing**

All proteins were placed in an octahedral solvation box with a 10 Å buffer between the protein and the nearest edge of the box. The system was treated using periodic boundary conditions. Approximately 8800 TIP3P water molecules were added to the solvation box. Sodium and Chloride ions were added to the box to balance the charge of the system. A table showing the counts of counter ions is provided below:

| **Protein Name** | **Sodium Ions** | **Chloride Ions** |
| --- | --- | --- |
| 6eqe_PET-tetramer_1 | 0 | 5 |
| 6eqe_1 | 0 | 5 |

**Forcefield Information**

All protein residues were parameterised using the AMBER ff19SB forcefield [Ref. [ff19SB](https://doi.org/10.1021/acs.jctc.9b00591)]. These parameters were prepared using tleap from the Ambertools package [Ref. [ambertools](https://doi.org/10.1021/acs.jcim.3c01153)]. Simulations were performed in explicit solvent. All water molecules parameterised using the TIP3P model [Ref. [tip3pParams](https://doi.org/10.1063/1.472061)]. Any ions in our system were treated using parameters calculated to complement the TIP3P water model [Ref. [ionParams(1)](https://doi.org/10.1021/ct500918t" \o "https://doi.org/10.1021/ct500918t), and [ionParams(2)](https://doi.org/10.1021/ct400146w" \o "https://doi.org/10.1021/ct400146w)].

**Simulation Protocols**

Initially, an energy minimisation step was performed using the steepest descent method. This energy minimisation step was performed until it reached convergence. A position restraint with a force constant of 500 kJ mol^-1^ nm^-2^ was applied to all atoms in the system.

Next, an energy minimisation step was performed using the steepest descent method. This energy minimisation step was performed until it reached convergence. A position restraint with a force constant of 430 kJ mol^-1^ nm^-2^ was applied to all atoms in the system.

Next, an energy minimisation step was performed using the steepest descent method. This energy minimisation step was performed until it reached convergence. A position restraint with a force constant of 360 kJ mol^-1^ nm^-2^ was applied to all atoms in the system.

Next, an energy minimisation step was performed using the steepest descent method. This energy minimisation step was performed until it reached convergence. A position restraint with a force constant of 290 kJ mol^-1^ nm^-2^ was applied to all atoms in the system.

Next, an energy minimisation step was performed using the steepest descent method. This energy minimisation step was performed until it reached convergence. A position restraint with a force constant of 220 kJ mol^-1^ nm^-2^ was applied to all atoms in the system.

Next, an energy minimisation step was performed using the steepest descent method. This energy minimisation step was performed until it reached convergence. A position restraint with a force constant of 150 kJ mol^-1^ nm^-2^ was applied to all atoms in the system.

Next, an energy minimisation step was performed using the steepest descent method. This energy minimisation step was performed until it reached convergence. A position restraint with a force constant of 80 kJ mol^-1^ nm^-2^ was applied to all atoms in the system.

Next, an energy minimisation step was performed using the steepest descent method. This energy minimisation step was performed until it reached convergence.

Next, a simulation was performed using the canonical (NVT) ensemble. This simulation was performed for 20 ps. The temperature of this simulation was stepped through the range 10 K, 20 K, 30 K, 40 K, 50 K, 60 K, 70 K, 80 K, and 100 K in even time increments. This simulation was performed using a mass of 4.03036 amu for hydrogen atoms. The mass added to each hydrogen atom was subtracted from the mass of the heavy atom it was bonded to. This, combined with the constraints placed upon bonds between heavy and hydrogen atoms, allowed the simulation to be performed using a timestep of 4 fs [Ref. [heavyProtons](https://doi.org/10.1002/(SICI)1096-987X(199906)20:8%3C786::AID-JCC5%3E3.0.CO;2-B" \o "https://doi.org/10.1002/(SICI)1096-987X(199906)20:8%3C786::AID-JCC5%3E3.0.CO;2-B)]. A position restraint with a force constant of 150 kJ mol^-1^ nm^-2^ was applied to all ligand atoms in the system.

Next, a simulation was performed using the *isothermal-isobaric* (NpT) ensemble. This simulation was performed for 9 ns. The temperature of this simulation was stepped through the range 100 K, 125 K, 150 K, 175 K, 200 K, 225 K, 250 K, 275 K, and 300 K in even time increments. This simulation was performed using a mass of 4.03036 amu for hydrogen atoms. The mass added to each hydrogen atom was subtracted from the mass of the heavy atom it was bonded to. This, combined with the constraints placed upon bonds between heavy and hydrogen atoms, allowed the simulation to be performed using a timestep of 4 fs [Ref. [heavyProtons](https://doi.org/10.1002/(SICI)1096-987X(199906)20:8%3C786::AID-JCC5%3E3.0.CO;2-B" \o "https://doi.org/10.1002/(SICI)1096-987X(199906)20:8%3C786::AID-JCC5%3E3.0.CO;2-B)]. A position restraint with a force constant of 150 kJ mol^-1^ nm^-2^ was applied to all ligand atoms in the system.

Next, a simulation was performed using the *isothermal-isobaric* (NpT) ensemble. This simulation was performed for 5 ns at 300 K. This simulation was performed using a mass of 4.03036 amu for hydrogen atoms. The mass added to each hydrogen atom was subtracted from the mass of the heavy atom it was bonded to. This, combined with the constraints placed upon bonds between heavy and hydrogen atoms, allowed the simulation to be performed using a timestep of 4 fs [Ref. [heavyProtons](https://doi.org/10.1002/(SICI)1096-987X(199906)20:8%3C786::AID-JCC5%3E3.0.CO;2-B" \o "https://doi.org/10.1002/(SICI)1096-987X(199906)20:8%3C786::AID-JCC5%3E3.0.CO;2-B)]. A position restraint with a force constant of 150 kJ mol^-1^ nm^-2^ was applied to all ligand atoms in the system.

Finally, a simulation was performed using the *isothermal-isobaric* (NpT) ensemble. This simulation was performed for 500 ns at 300 K. This simulation was performed using a mass of 4.03036 amu for hydrogen atoms. The mass added to each hydrogen atom was subtracted from the mass of the heavy atom it was bonded to. This, combined with the constraints placed upon bonds between heavy and hydrogen atoms, allowed the simulation to be performed using a timestep of 4 fs [Ref. [heavyProtons](https://doi.org/10.1002/(SICI)1096-987X(199906)20:8%3C786::AID-JCC5%3E3.0.CO;2-B)].

All simulations were performed using the OpenMM simulation toolkit [Ref. [openmm](https://doi.org/10.1371/journal.pcbi.1005659" \o "https://doi.org/10.1371/journal.pcbi.1005659)]. All simulations were performed using the Langevin Middle Integrator [Ref. [langevinMiddleIntegrator](https://doi.org/10.1021/acs.jpca.9b02771" \o "https://doi.org/10.1021/acs.jpca.9b02771)] which was used to enforce constant temperature conditions in each simulation. For simulations run under the *isothermal-isobaric* (NpT) ensemble, the Monte-Carlo barostat was used to enforce a constant pressure of 1 bar. In all simulations, long-range Coulombic interactions were modelled using the Particle-Mesh Ewald (PME) method [Ref. [pme](https://doi.org/10.1063/1.464397" \o "https://doi.org/10.1063/1.464397)], with a 10 Å cutoff distance. In all simulations, constraints were applied to bonds between hydrogen atoms and heavy atoms using the SHAKE algorithm [Ref. [shake](https://doi.org/10.1002/1096-987X(20010415)22:5%3C501::AID-JCC1021%3E3.0.CO;2-V)]. In all simulations, bonds lengths and angles of water molecules were constrained using the SETTLE algorithm [Ref. [settle](https://doi.org/10.1002/jcc.540130805)].

#### Human-written methods section for molecular dynamics protocol

##### Parameterisation

To the AMBER ff19SB forcefield [7] was used to parameterise the protein portion of our system. We use the TIP3P parameter set [8] to describe the water molecules in our system. Ions in our system were parameterised using AMBER parameters that are optimized for interactions with TIP3P waters [9]. To generate parameters for our PET-tetramer ligand, we first calculated partial charges using antechamber [9,10], then generated AMBER compatible parameter files using parmchk [9,10]. We finally generated our system parameter files using TLEAP from the ambertools package [11].

##### Solvation, Equilibration, and Simulation

Our systems were placed in a cubic water box with a 10 Å buffer between our protein and the edges of the solvent box. TIP3P [8] water molecules in this box were replaced with chloride ions to ensure the overall charge of the system was neutralized.

A series of seven steepest-decent energy minimisation steps were then performed on each system. Each energy minimisation step was performed with position restraints placed on all atoms. The force constant for these restraints in the first simulation was 500 kJ mol^-1^ Å^—2^. In each subsequent energy minimisation step, we reduced the force constant by 70 kJ mol^-1^ Å^--2^. In the last of these energy minimisation steps, no position restraints were used. Each of these energy minimisation steps were run until they reached convergence.

We then performed a 20 ps MD simulation under the *canonical* ensemble (NVT / constant volume). During this NVT simulation we heated the system from 0 K to 100 K in 9 steps of roughly 2.2 ps each. A position restraint was applied to all atoms in our PET-tetramer ligand with a force constant of 150 kJ mol^-1^. We then performed a 9 ns MD simulation under the *isothermal-isobaric* ensemble (NpT / constant pressure). During this NpT simulation, we heated the system from 100 K to 300 K in 9 steps of 1 ns each. A position restraint was applied to all atoms in our PET-tetramer ligand with a force constant of 150 kJ mol^-1^. We the performed a further 5 ns MD simulation at 300 K under NpT conditions to fully equilibrate our systems. A position restraint was applied to all atoms in our PET-tetramer ligand with a force constant of 150 kJ mol^-1^. Finally, production MD simulations were performed for 500 ns at 300K under NpT conditions. No restraints were applied in our production MD simulations.

All simulations were performed using the OpenMM simulation toolkit [1]. The following methods were used in all MD simulations: Velocities of the system were calculated using the Langevin middle integrator [12]. Non-bonded interactions were modelled using the Particle-Mesh Ewald method with a non-bonded cutoff of 10 Å [13]. Constraints were applied to all bonds between heavy atoms and hydrogen atoms using the SHAKE algorithm [14]. Bonds and angles of water molecules were restrained using the SETTLE algorithm [15]. For NpT simulations, the Monte-Carlo barostat was used to enforce a constant pressure of 1 bar. We treated all protons as having a mass of 4.03036 amu, the extra mass being subtracted from adjacent heavy atoms [16]. This, combined with hydrogen bond constraints allowed all simulations to be performed using a timestep of 4 fs, which roughly doubled our simulation speed.

All the above information is stored in the config file used to run all of these simulations.

All simulations were performed in an Alienware m16 R1 laptop with an intel i9 processor and NVIDIA GeForce RTX 4080 graphics card.

##### Analysis of trajectories

All analyses were performed using a the MDAnalysis software package [2,3]. Plots were made using the Matplotlib python package. The scripts for running these are freely available on our GitHub: <https://github.com/wells-wood-research/drMD>

#### Configuration file

The following file (located in the Prescriptions directory in the drMD repository) PETase_MD_config.yaml was used to perform simulations of PETase with and without a PET-tetramer ligand.

```
########################################################################
pathInfo:
  inputDir: /home/esp/scriptDevelopment/drMD/01_inputs/PET_proj
  outputDir: /home/esp/scriptDevelopment/04_PET_proj_outputs
########################################################################
miscInfo:
  pH: 7.4
  firstAidMaxRetries: 10
  boxGeometry: octahedral
  skipPdbTriage: false
  firstAidMaxRetries: 100
  writeMyMethodsSection: True
  trajectorySelections:
  - selection:
      keyword: protein
  - selection:
      keyword: ligand
########################################################################
hardwareInfo:
  parallelCPU: 1
  platform: CUDA
  subprocessCpus: 1
########################################################################
simulationInfo:
## EM STEPS ##################################
  - stepName: 011_energy_minimisation
    simulationType: EM
    temperature: 300
    maxIterations: 1000
    restraintInfo:
    - restraintType: position
      parameters:
        k: 500
      selection:
        keyword: all

  ##
  - stepName: 012_energy_minimisation
    simulationType: EM
    temperature: 300
    maxIterations: 10000
    restraintInfo:
    - restraintType: position
      parameters:
        k: 430
      selection:
        keyword: all

  ##
  - stepName: 013_energy_minimisation
    simulationType: EM
    temperature: 300
    maxIterations: 10000
    restraintInfo:
    - restraintType: position
      parameters:
        k: 360
      selection:
        keyword: all

  ##
  - stepName: 014_energy_minimisation
    simulationType: EM
    temperature: 300
    maxIterations: 10000
    restraintInfo:
    - restraintType: position
      parameters:
        k: 290
      selection:
        keyword: all

  ##
  - stepName: 015_energy_minimisation
    simulationType: EM
    temperature: 300
    maxIterations: 10000
    restraintInfo:
    - restraintType: position
      parameters:
        k: 220
      selection:
        keyword: all

  ##
  - stepName: 015_energy_minimisation
    simulationType: EM
    temperature: 300
    maxIterations: 10000
    restraintInfo:
    - restraintType: position
      parameters:
        k: 150
      selection:
        keyword: all

  ##
  - stepName: 016_energy_minimisation
    simulationType: EM
    temperature: 300
    maxIterations: 10000
    restraintInfo:
    - restraintType: position
      parameters:
        k: 80
      selection:
        keyword: all

  ##
  - stepName: 017_energy_minimisation
    simulationType: EM
    temperature: 300
    maxIterations: 10000
## EQUILIBRATION STEPS #######################
  - stepName: 021_NVT_warmup
    simulationType: NVT
    duration: 20 ps
    heavyProtons: true
    timestep: 4 fs
    temperatureRange: [10,20,30,40,50,60,70,80,100]
    logInterval: 1 ps
    restraintInfo:
    - restraintType: position
      parameters:
        k: 150
      selection:
        keyword: ligand
  ##
  - stepName: 022_NpT_warmup
    simulationType: NPT
    duration: 9 ns
    heavyProtons: true
    timestep: 4 fs
    temperatureRange: [100, 125, 150, 175, 200, 225, 250, 275, 300]
    logInterval: 100 ps
    restraintInfo:
    - restraintType: position
      parameters:
        k: 150
      selection:
        keyword: ligand

  ##
  - stepName: 023_NpT_equilibration
    simulationType: NPT
    duration: 5 ns
    heavyProtons: true
    timestep: 4 fs
    temperature: 300
    logInterval: 100 ps
    restraintInfo:
    - restraintType: position
      parameters:
        k: 150
      selection:
        keyword: ligand
## PRODUCTION MD #######################
  - stepName: 031_production_md
    simulationType: NPT
    duration: 500 ns
    heavyProtons: true
    timestep: 4 fs
    temperature: 300
    logInterval: 10 ps
```

#### Simulation Health Checks

This section contains snapshots of the simulation “vitals” report generated at the end of the equilibration and production MD steps for systems I and II.

##### Vitals reports for system I

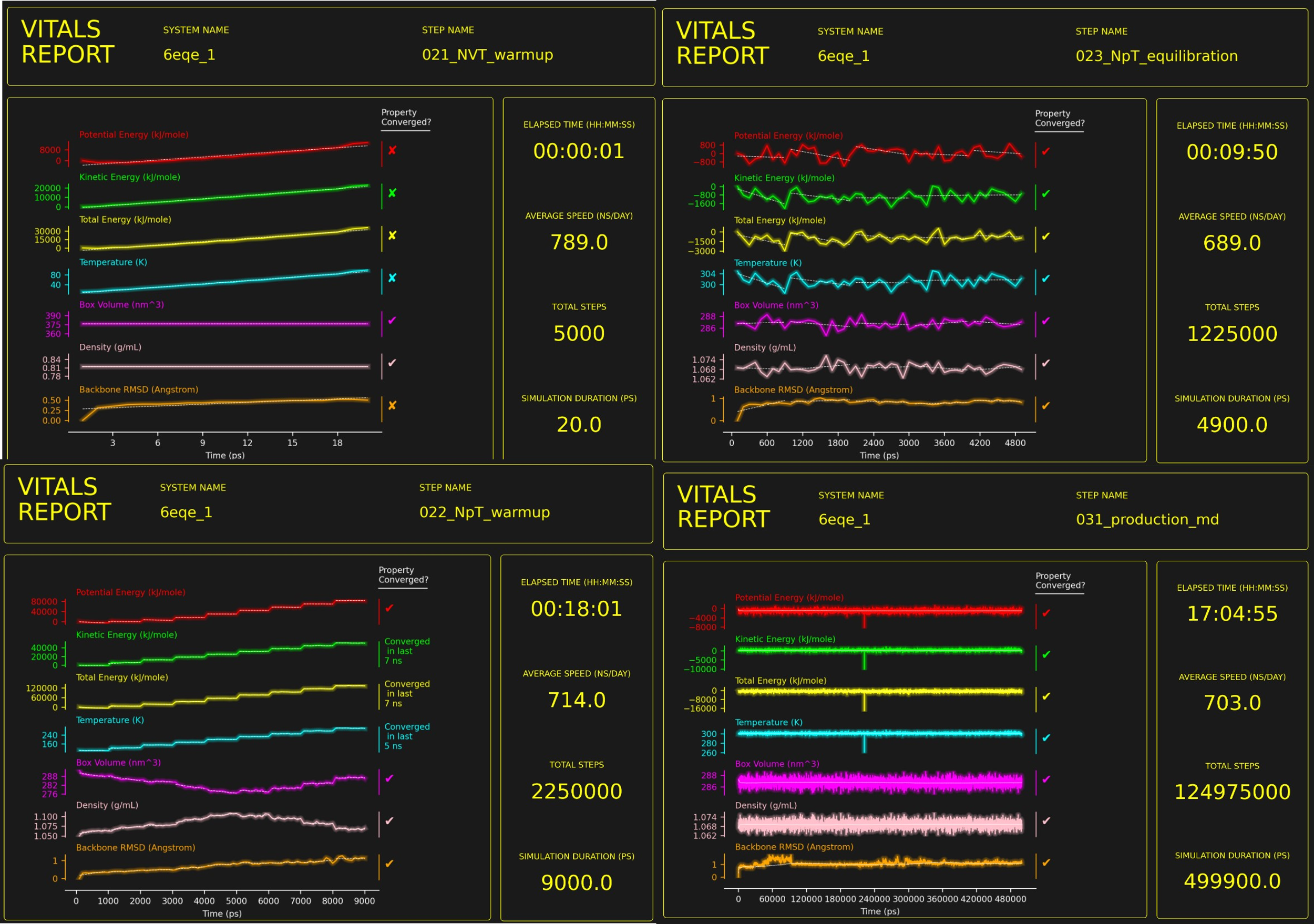

Figure SI III: Vitals reports for system I (Apo isPETase) for all four simulation steps of our protocol.

##### Vitals reports for system II

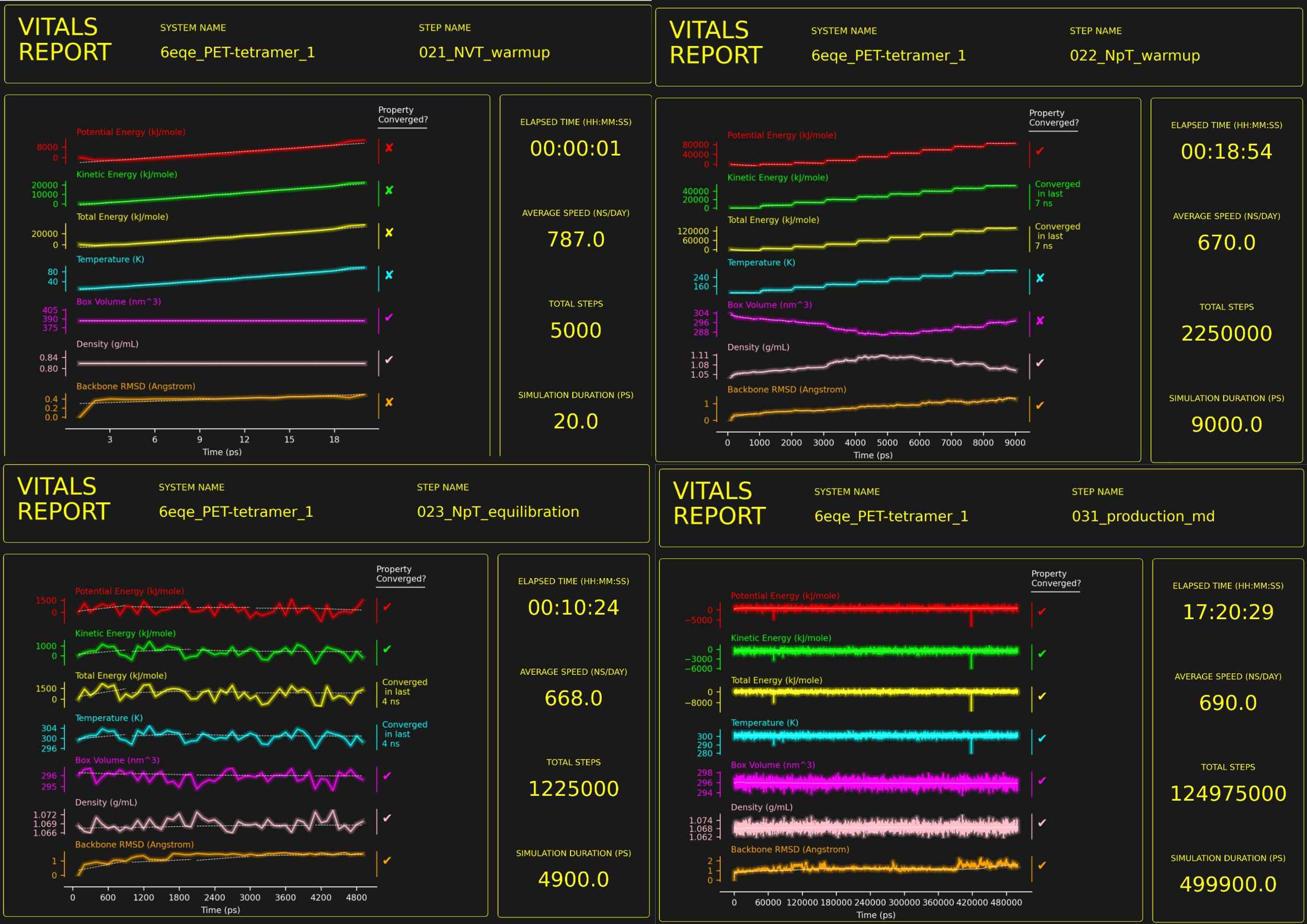

Figure SI IV: Vitals reports for system II (Holo isPETase + PET -tetramer) for all four simulation steps of our protocol.

##### Example interpretation of vitals reports

For the two systems above, we have performed four simulation steps (after the seven energy minimisation steps, which are not considered simulation steps). After each simulation step is completed, drMD will generate a PDF file containing a vitals report for the simulation.

Contained in each vitals report the following information is presented: System and step names across the top and time-related data on the right. In the main panel of the report, traces for the following properties of the simulation are plotted vs simulation time:

- Potential Energy
- Kinetic Energy
- Total Energy
- Temperature
- Simulation box volume
- Simulation box density
- Backbone RMSD

Typically, the question a researcher wants to answer immediately after running a simulation is: Has my simulation converged? This is important as a converged simulation can be sampling from equilibrium conditions. As most real-world systems operate at equilibrium conditions, measurements from simulations can only be considered valid if the simulation is at equilibrium too. Traditionally, this is determined by looking at a plot of the system’s RMSD and determining if the trace is “flat”.

If we look at the “021_NVT_warmup” step for either system, we can see the traces for all three energy properties as well as temperature steadily rising throughout the simulation, whilst the box volume and density remain constant (as expected for an N**V**T simulation). For the “022_NpT_warmup”, we can see that all three energy properties as well as temperature rise in distinct steps, while the box volume and density fluctuate. From these traces, we can first determine that neither of these simulations are at equilibrium yet. We can also tell that these simulations are serving their purpose to gradually heat up our system to the desired temperature of 300 K.

If we then look at the “023_NpT_Equilibiation” step for either system, we can see that after a few nanoseconds of simulation time, the RMSD of the system converges, showing that our system is now sampling from equilibrium conditions. We can now have some confidence that measurements from this point onwards will be representative of a “real” system at equilibrium. If we compare the traces of the three energy properties for the two systems, we can see that the traces for the Apo (system I) converge while significant fluctuations can be observed for Holo (system II). This is most likely due to the presence of the highly flexible PET-tetramer in system II.

Next, if we look at the reports for “031_production_MD” for each system, we can see that all properties show mostly “flat” traces. This is good as it further supports the notion that we are sampling from an equilibrated simulation. By looking at the RMSD trace, we can identify “events” in the simulation when the RMSD value spikes, then returns to a more constant state. These “events” could represent large-scale motions of the protein backbone and may warrant further investigation.
